# Semaphorin Receptors Antagonize Wnt Signaling Through Beta-Catenin Degradation

**DOI:** 10.1101/2024.05.29.596372

**Authors:** Tyler M. Hoard, Katie Liu, Kenneth M. Cadigan, Roman J. Giger, Benjamin L. Allen

## Abstract

Precise control of morphogen signaling levels is essential for proper development. An outstanding question is: what mechanisms ensure proper morphogen activity and correct cellular responses? Previous work has identified Semaphorin (SEMA) receptors, Neuropilins (NRPs) and Plexins (PLXNs), as positive regulators of the Hedgehog (HH) signaling pathway. Here, we provide evidence that NRPs and PLXNs antagonize Wnt signaling in both fibroblasts and epithelial cells. Further, *Nrp1/2* deletion in fibroblasts results in elevated baseline Wnt pathway activity and increased maximal responses to Wnt stimulation. Notably, and in contrast to HH signaling, SEMA receptor-mediated Wnt antagonism is independent of primary cilia. Mechanistically, PLXNs and NRPs act downstream of Dishevelled (DVL) to destabilize β-catenin (CTNNB1) in a proteosome-dependent manner. Further, NRPs, but not PLXNs, act in a GSK3b/CK1-dependent fashion to antagonize Wnt signaling, suggesting distinct repressive mechanisms for these SEMA receptors. Overall, this study identifies SEMA receptors as novel Wnt pathway antagonists that may also play larger roles integrating signals from multiple inputs.

## Introduction

Semaphorin (SEMA) signaling is an important regulator of cell migration, axon guidance, immune system function, angiogenesis, and tissue repair (Jongbloets and Pasterkamp 2014; Koropouli and Kolodkin 2014; Fard and Tamagnone 2021). Neuropilin (NRP) and Plexin (PLXN) family members function as single-pass transmembrane co-receptors for class 3 SEMA proteins (Chen et al. 1997; He and Tessier-Lavigne 1997; Kolodkin et al. 1997; Takahashi et al. 1999; Tamagnone et al. 1999; Gu et al. 2005). NRPs consist of 2 members (NRP1 and NRP2) while PLXNs consist of 9 members divided into 4 subfamilies based on structural and functional differences and similarities (PLXNA1-4, PLXNB1-3, PLXNC1, and PLXND1) (Pellet-Many et al. 2008; Perala et al. 2012). Upon SEMA binding to the NRP and PLXN co-receptor complex, catalytic activity of the intracellular PLXN bipartite GTPase-activating Protein (GAP) domain results in cytoskeletal and cell adhesion changes (Perala et al. 2012).

In addition to their roles in SEMA signaling, PLXNs and NRPs act promiscuously to affect multiple other pathways, including VEGF signaling and, more recently, the Hedgehog (HH) pathway (Soker et al. 1998; Kawasaki et al. 1999; Fukushima et al. 2011; Hillman et al. 2011; Kim et al. 2011; Ge et al. 2015; Pinskey et al. 2017; Pinskey et al. 2022). These data raise the question of whether PLXNs and NRPs may regulate other key developmental signaling pathways.

Wnt signaling is a highly conserved pathway that plays important roles in proper cell differentiation, organogenesis, tissue maintenance and repair, and adult stem cell function; conversely, aberrant Wnt signaling can drive pathogenesis (Clevers and Nusse 2012; Nusse and Clevers 2017; Wiese et al. 2018). The Wnt pathway consists of several core components, including the glycosylated Wnt ligands, the seven-pass transmembrane receptor, Frizzled (FZD), the single pass transmembrane protein co-receptors, Low Density Lipoprotein Receptor-Related Protein (LRP5/6), transcriptional co-activator, β-catenin (CTNNB1), and transcription factors belonging to the T-Cell Factor/Lymphoid Enhancer Factor (TCF/LEF) family (McCrea and Crute 1991; Behrens et al. 1996; Molenaar et al. 1996; Yang-Snyder et al. 1996; Yamamoto et al. 1998; Wehrli et al. 2000; Willert et al. 2003). In the absence of Wnt ligand, CTNNB1 is phosphorylated, ultimately leading to its ubiquitination and subsequent proteolytically degradation. These phosphorylation events take place at four conserved residues within the N-terminal region of CTNNB1 (Stamos and Weis 2013). S45 is phosphorylated by Casein Kinase 1 (CK1), a permissive step that allows for the subsequent phosphorylation of T41, S37, and S33 by Glycogen Synthase 3 β (GSK3β) (Stamos and Weis 2013). CTNNB1 phosphorylation takes place amongst a group of associated proteins in the cytoplasm including the scaffolding proteins, AXIN2 and APC (Stamos and Weis 2013). Upon Wnt ligand binding to FZD, Disheveled (DVL) is phosphorylated, resulting in recruitment of AXIN2 and GSK3β to the cell membrane, preventing the proteasomal turnover of CTNNB1. This allows CTNNB1 to translocate to the nucleus and function in tandem with TCF/LEF to activate Wnt targets (Smalley et al. 1999; Stamos and Weis 2013).

In this study, we investigated the role of PLXNs and NRPs in Wnt signaling. Our data indicate that expression of either PLXNs or NRPs antagonize Wnt signaling in both NIH/3T3 fibroblasts and HEK293T epithelial cells. Further, genetic deletion of *Nrp1* and *Nrp2* in NIH/3T3 fibroblasts results in increased baseline Wnt pathway activity and a greater maximal response to CHIRON-mediated Wnt pathway activation. Using these cells, we also find the PLXNs antagonize Wnt signaling in a NRP-independent fashion. Notably, and in contrast to HH signaling, PLXN/NRP-mediated Wnt antagonism is independent of the primary cilium, a key organelle that has been previously implicated in Wnt signaling. Using triple knockout cells, we find that SEMA receptor Wnt pathway antagonism is also DVL-independent. Instead, PLXNs and NRPs restrict Wnt signaling through proteasome-mediated CTNNB1 degradation. Notably, while mutagenesis studies indicated that NRP-mediated CTNNB1 degradation is GSK3B/CK1-dependent, PLXN-mediated CTNNB1 degradation acts through a distinct, albeit yet to be determined mechanism. Together, these data identify SEMA receptors as novel Wnt pathway antagonists.

## Results

### PLXNs and NRPs antagonize Wnt signaling in NIH/3T3 fibroblasts

In a previous study defining PLXNs as novel positive regulators of the HH signaling pathway, we preliminarily observed that PLXNs antagonize Wnt signaling (Pinskey et al. 2022). To further investigate the contributions of SEMA receptors to Wnt and HH signaling (Figure 1), we initially confirmed our previous results using HH-responsive and Wnt-responsive luciferase assays in NIH/3T3 fibroblasts (Figure 2A,B). HH pathway activation via expression of an oncogenic Smo construct (*SmoM2*) is enhanced by *Plxna1* co-expression (Figure 2A), while Wnt pathway stimulation via expression of a stabilized *CTNNB1* construct (*CTNNB1^S33Y^*) is inhibited by *Plxna1* co-expression (Figure 2B). In addition to *Plxna1*, all members of the *Plxna* subfamily as well as *Plxnb2* antagonize Wnt pathway activity (Figure S1 A-D). To control for potential non-specific effects of cell surface receptor expression on Wnt signaling, we expressed *Ptch1*, the canonical HH pathway receptor, which does not antagonize Wnt signaling in this assay (Figure S1E).

**Figure 1.**
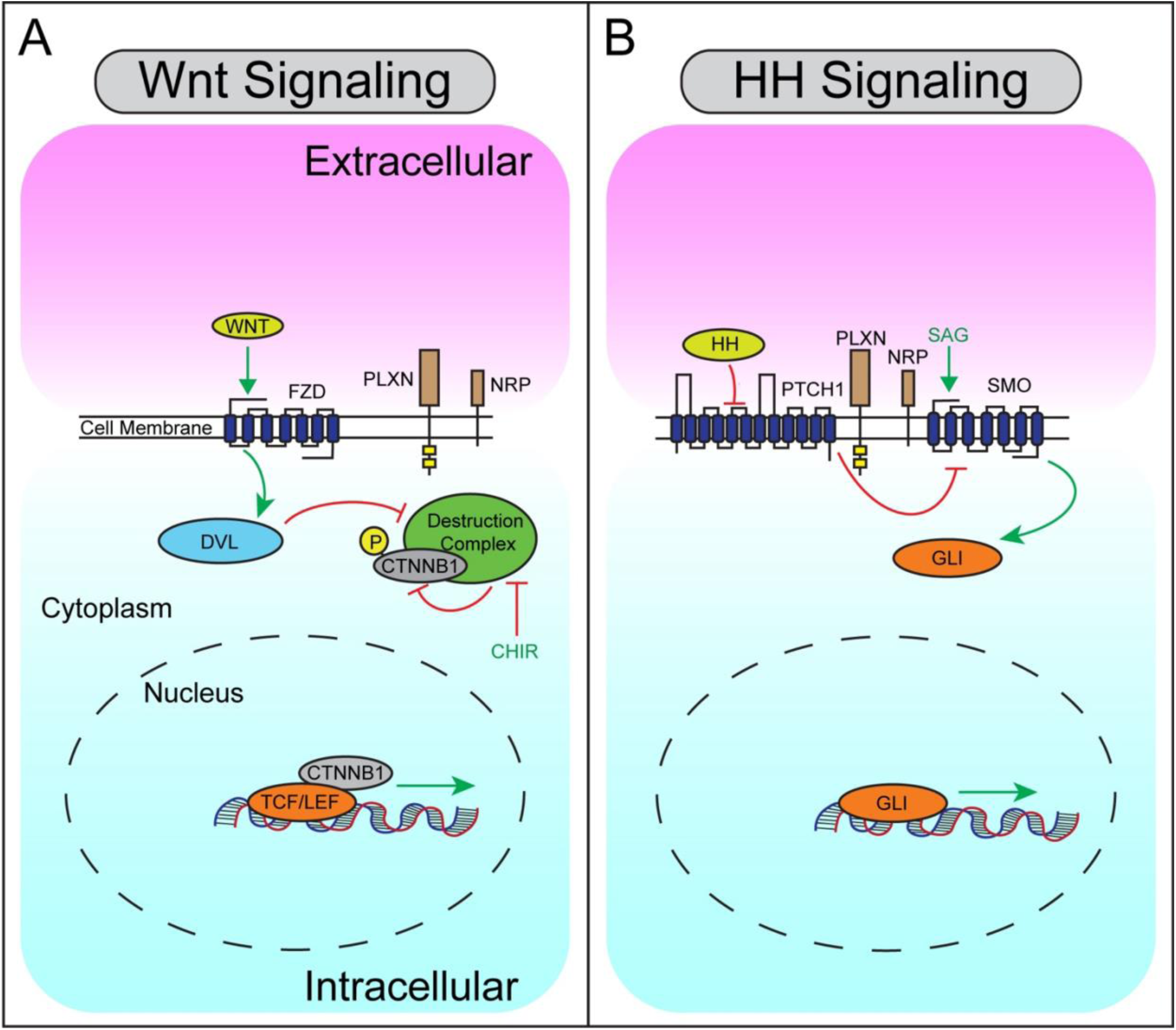
Simplified Schematic of Wnt and HH Signaling Pathways. (**A**) Summary of Wnt signaling in a Wnt-responsive cell (extracellular, pink; intracellular, blue). Wnt ligands initiate signaling through the binding and activation of FZD receptors, which act through DVL to inhibit the destruction complex (consisting of Axin, APC, GSK3β, and CK1α), preventing CTNNB1 phosphorylation, leading to CTNNB1 stabilization in the cytoplasm and nuclear translocation. In the nucleus, CTNNB1 interacts with TCF/LEF transcription factors to activate Wnt target genes. CHIR (green) is a small molecule GSK3β antagonist that activates the Wnt pathway. (**B**) Summary of HH signaling in a HH-responsive cell. HH ligands bind to the canonical receptor PTCH1, inhibiting its activity and leading to de-repression of SMO, initiating a signal transduction cascade that culminates in GLI transcription factor translocation to the nucleus and activation of HH target genes. SAG (green) is a small molecule SMO agonist that activates the HH pathway. PLXNs and NRPs (brown) are expressed at the cell surface in both Wnt- and HH-responsive cells.

**Figure 2.**
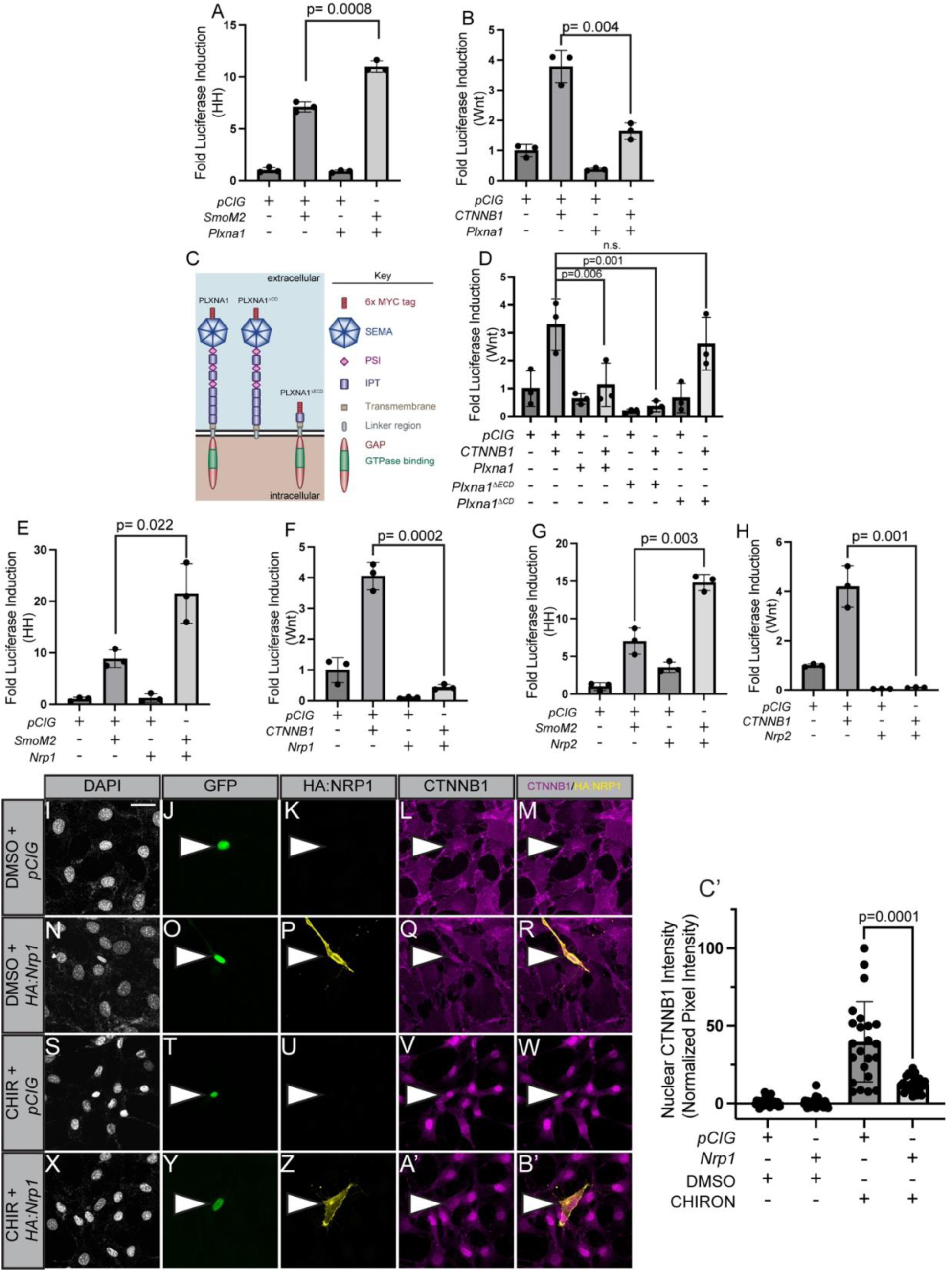
PLXN or NRP expression in NIH/3T3 fibroblasts promotes HH signaling and antagonizes Wnt signaling. (**A-B**) HH- or Wnt-dependent luciferase reporter activity (indicated on the y-axis) was measured in NIH/3T3 cells transfected with the indicated plasmids (*pCIG* = empty vector). (**C**) Schematic representation of different PLXNA1 proteins. (**D-H**) HH- or Wnt-dependent luciferase reporter activity was measured in NIH/3T3 cells transfected with the indicated plasmids. (**I-B’**) NIH/3T3 cells were transfected with the indicated plasmids and treated with either CHIRON or vehicle (DMSO). Antibodies were used to detect HA:NRP1 (yellow) and CTNNB1 (magenta). Nuclear GFP (green) indicates transfected cells while DAPI (gray) stains all nuclei. Scale bar = 10 μm. White arrowheads denote transfected cells. (**C’**) Quantitation of nuclear CTNNB1 in **I-B’**. Pixel intensity of nuclear CTNNB1 staining was measured and normalized to cytoplasmic pixel intensity. Data points indicate technical replicates. Data are representative of at least three biological replicates. Data are reported as mean fold change +/- S.D., with p-values calculated using two-tailed unpaired Student’s t-test in **A-B, E-H,** and **C’** and a one-way ANOVA used in **D**. n.s., not significant.

To determine the domains necessary for PLXN-mediated Wnt pathway antagonism, we utilized constructs lacking either the PLXN extracellular (*Plxna1*^Δ*ECD*^) or cytoplasmic domains (*Plxna1*^Δ*CD*^; Figure 2C). *Plxna1*^Δ*ECD*^ antagonizes Wnt signaling to an even greater degree than full-length *Plxna1* (Figure 2D), consistent with a previously described autoinhibitory function for the PLXN extracellular domain (Takahashi and Strittmatter 2001). Conversely, *Plxna1*^Δ*CD*^ expression does not alter Wnt signaling (Figure 2D). These data suggest that PLXN-mediated Wnt pathway antagonism requires an intact cytoplasmic domain.

Given that both PLXNs and NRPs promote HH signaling, we wondered whether PLXNs and NRPs also both antagonize Wnt signaling (Hillman et al. 2011; Ge et al. 2015; Pinskey et al. 2017; Pinskey et al. 2022). Interestingly, both *Nrp1* (Figure 2E-F) and *Nrp2* (Figure 2G-H) antagonize CTNNB1-mediated Wnt pathway activation. Notably, *Nrp1* or *Nrp2* expression restricts baseline Wnt pathway activity in NIH/3T3 fibroblasts (Figure 2F, H), raising the question of the effects of *Nrp* expression on endogenous CTNNB1. Following CHIRON-mediated stimulation of the Wnt pathway in NIH/3T3 fibroblasts, we observed a significant decrease in nuclear endogenous nuclear CTNNB1 in *Nrp1*-expressing compared to control transfected GFP-expressing cells (*pCIG)* (Figure 2I-C’). These data indicate that both PLXNS and NRPs can antagonize Wnt pathway activity.

### *Nrp1* and *Nrp2* deletion results in elevated baseline Wnt pathway activity

Since *Nrp* expression abrogated baseline Wnt pathway activity (Figure 2F, H) and reduced endogenous CTNNB1 nuclear localization (Figure 2C’), we investigated the consequences of *Nrp* deletion on baseline Wnt signaling. We utilized CRISPR-mediated gene editing to delete both *Nrp1* and *Nrp2* in NIH/3T3 cells (Figure 3A, Figure S2A-F). We validated loss of NRP1 and NRP2 by Western Blot analysis (Figure 3B). Additionally, we assessed the HH responsiveness of *Nrp1^-/-^;Nrp2^-/-^* cells following stimulation by Smoothened agonist (SAG) treatment (Figure 3C). Consistent with previous reports, combined *Nrp1/2* deletion results in a loss of HH pathway responsiveness (Figure 3C), while re-expression of *Nrp1* and *Nrp2* restores SAG-mediated HH signaling in these cells (Figure S2G) (Hillman et al. 2011; Ge et al. 2015).

**Figure 3.**
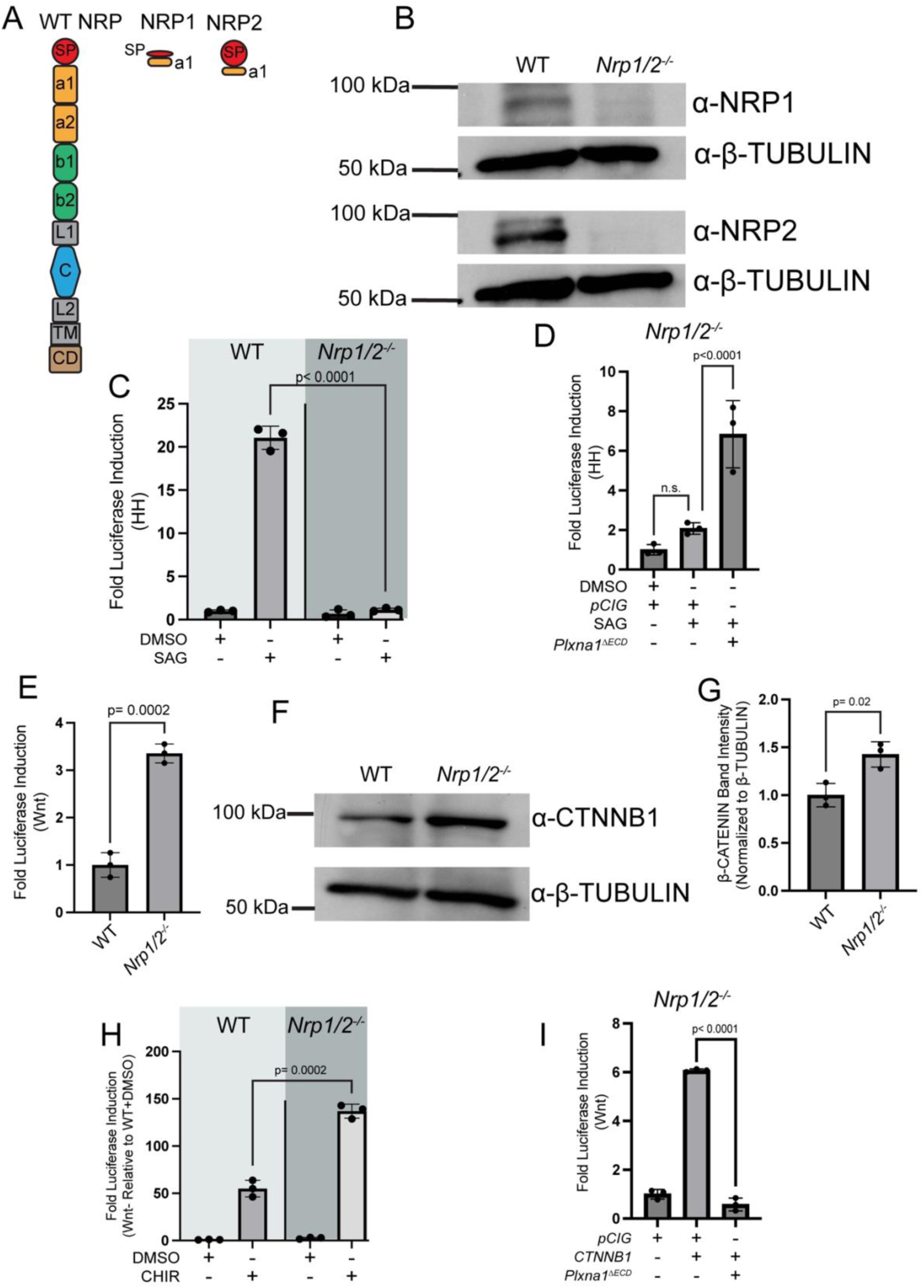
*Nrp1^-/-^Nrp2^-/-^*NIH/3T3 cells display elevated baseline Wnt signaling. (**A**) Schematic representation of wildtype (WT) and CRISPR-edited forms of NRP1 and NRP2. (**B**) Western blot analysis confirmed deletion of NRP1 and NRP2 in NIH/3T3 cells. Anti-β-TUBULIN was used as a loading control. (**C**) WT or *Nrp1^-/-^Nrp2^-/-^* NIH/3T3 cells were treated with DMSO or SAG and HH pathway activity was measured by luciferase assay. (**D**) *Nrp1^-/-^ Nrp2^-/-^* NIH/3T3 cells were transfected with empty vector (*pCIG*) or *Plxna1*^Δ*ECD*^ and treated with DMSO or SAG; HH signaling levels were measured by luciferase assay. (**E**) Baseline Wnt-dependent luciferase reporter activity was measured in untreated WT and *Nrp1^-/-^;Nrp2^-/-^*NIH/3T3 cells transfected with the TOPFlash Wnt reporter. (**F**) Western blot analysis analyzing levels of CTNNB1 in WT and *Nrp1^-/-^;Nrp2^-/-^*NIH/3T3 cells. Anti-β-TUBULIN was used as a loading control. (**G**) Quantitation of CTNNB1 intensity in (**C**) normalized to β-TUBULIN intensity. (**H**) WT or *Nrp1^-/-^Nrp2^-/-^* NIH/3T3 cells were treated with DMSO or CHIRON, and Wnt signaling levels were measured by luciferase assay. Data are represented relative to the first condition (WT cells + DMSO). (**I**) *Nrp1^-/-^Nrp2^-/-^* NIH/3T3 cells were transfected with the indicated plasmids and Wnt signaling levels were measured by luciferase assay. Data are representative of at least three biological replicates. Data are reported as mean fold change +/- S.D., with p-values calculated using two-tailed unpaired Student’s t-test in **C, E,** and **G-I** and a one-way ANOVA in **D**. n.s., not significant.

Notably, *Nrp1^-/-^;Nrp2^-/-^* cells still robustly activate HH signaling following transfection with *Gli1* (Figure S2H-I), which encodes for a constitutive transcriptional activator of HH signaling. While PLXNs and NRPs promote GLI-mediated HH pathway activity, they are not required for GLI function (Hillman et al. 2011; Pinskey et al. 2022). Surprisingly, *Plxna1*^Δ*ECD*^ expression results in HH pathway stimulation in *Nrp1^-/-^;Nrp2^-/-^* NIH/3T3 cells (Figure 3D), indicating that PLXN-mediated HH pathway promotion is NRP-independent.

Having validated efficient *Nrp1/2* deletion and confirmed predicted effects on HH signaling, we assessed baseline Wnt signaling in these cells, where we observed an approximate 3-fold increase in baseline Wnt pathway activity compared to WT NIH/3T3 cells (Figure 3E).

We also observed an increase in CTNNB1 by western blot analysis in *Nrp1^-/-^;Nrp2^-/-^* NIH/3T3 cells compared to WT NIH/3T3 cells (Figure 3F-G). Further, *Nrp1^-/-^;Nrp2^-/-^* cells display increased Wnt signaling compared to WT NIH/3T3 cells treated with the same concentration of CHIRON (Figure 3H). Together, these data suggest that *Nrp1/2* deletion results in increased Wnt baseline activity as well as higher Wnt pathway induction following CHIRON stimulation.

Given that *Plxna1*^Δ*ECD*^ promotes HH signaling in a NRP-independent fashion, we tested whether PLXN-mediated Wnt antagonism was also NRP-independent. Notably, *Plxna1*^Δ*ECD*^ expression results in near-complete Wnt pathway inhibition in *Nrp1^-/-^;Nrp2^-/-^*NIH/3T3 cells (Figure 3I). Importantly, re-expression of either *Nrp1* or *Nrp2* in *Nrp1/2* KO cells restricted Wnt pathway activity, although *Nrp2* appears to be more effective than *Nrp1* (Figure S2J). Together, these data suggest that PLXN-mediated HH pathway promotion and Wnt pathway inhibition are NRP-independent.

### PLXNs and NRPs antagonize Wnt signaling independently of primary cilia

We previously found that PLXNs require intact primary cilia to promote HH signaling (Pinskey et al. 2022). Consistent with our previous observations, SAG fails to activate HH signaling in *Kif3a^-/-^* NIH/3T3 cells, which lack primary cilia (Figure 4A-B). While HH signaling can be stimulated in *Kif3a^-/-^* cells following *Gli1* transfection, co-expression with *Plxna1*^Δ*ECD*^ fails to further promote HH signaling in *Kif3a^-/-^* cells compared to WT NIH/3T3 cells (Figure 4C-D).

**Figure 4.**
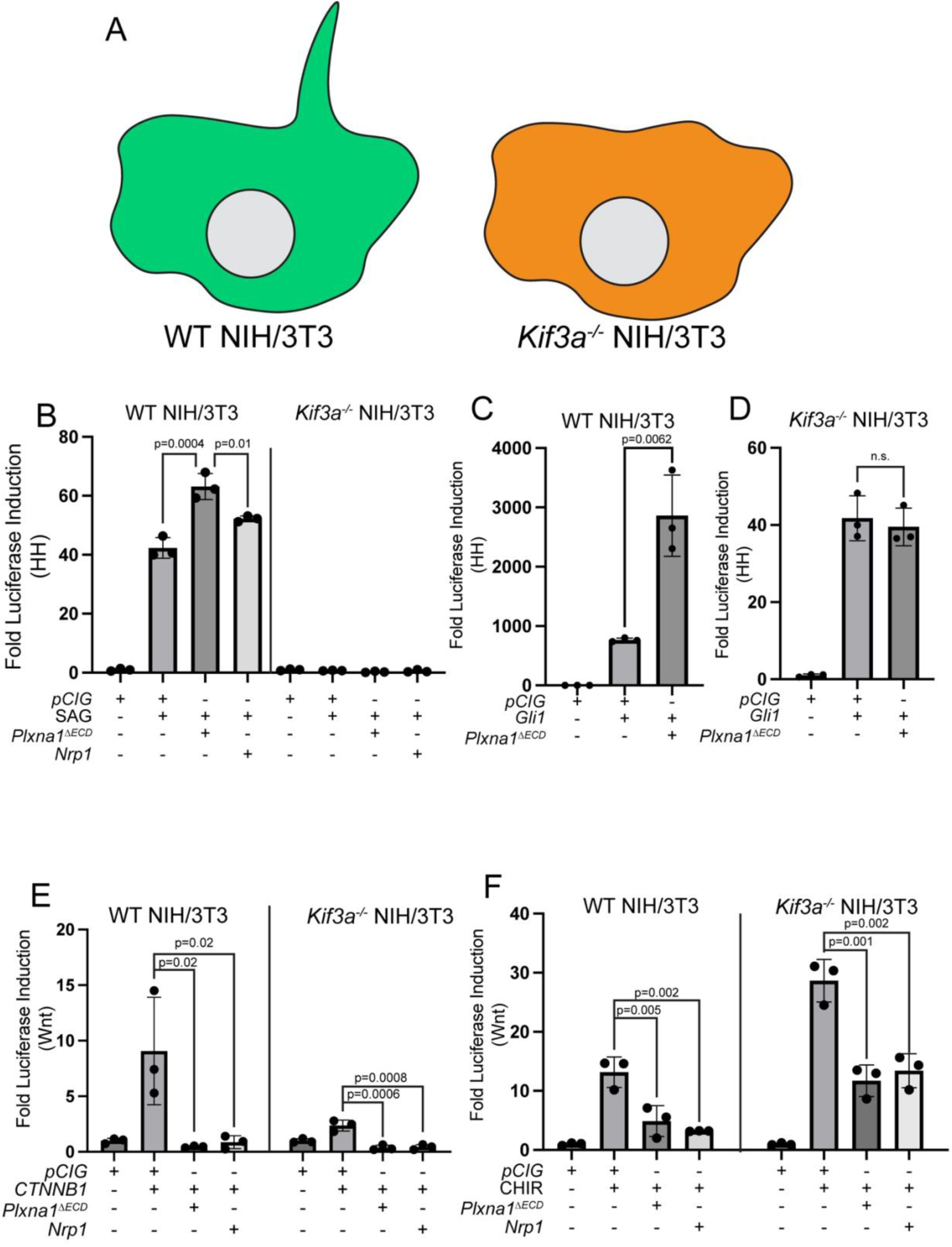
PLXN and NRP antagonize Wnt signaling independently of primary cilia. (**A**) Schematic of a WT NIH/3T3 cell containing a primary cilium and a *Kif3a^-/-^* NIH/3T3 cell, which lacks a primary cilium. (**B-D**) WT or *Kif3a^-/-^* NIH/3T3 cells were transfected with the indicated plasmids and HH signaling was measured by luciferase assay. (**E-F**) WT or *Kif3a^-/-^* NIH/3T3 cells were transfected with the indicated plasmids, and Wnt signaling was measured by luciferase assay. Data are representative of at least three biological replicates. Data are reported as mean fold change +/- S.D., with p-values calculated using a one-way ANOVA in **B, E**, and **F**, and a two-tailed unpaired Student’s t-test in **C** and **D**. n.s., not significant.

The role of primary cilia in Wnt signaling remains controversial (Vuong and Mlodzik 2023). We found that CTNNB1-mediated Wnt pathway activation is blunted in *Kif3a^-/-^* cells compared to WT NIH/3T3 cells (Figure 4E). However, both *Plxna1*^Δ*ECD*^ and *Nrp1* both still repress CTNNB1-mediated Wnt pathway activation in *Kif3a^-/-^* NIH/3T3 cells (Figure 4E).

Further, CHIRON-mediated Wnt pathway activation is similar between WT and *Kif3a^-/-^* NIH/3T3 cells (Figure 4F). Again, both *Plxna1*^Δ*ECD*^ and *Nrp1* antagonize CHIRON-mediated Wnt pathway activation in *Kif3a^-/-^* NIH/3T3 cells (Figure 4F). These data indicate that PLXNs and NRPs antagonize Wnt signaling in a primary cilia-independent fashion, identifying an important mechanistic divergence from PLXN/NRP-mediated promotion of HH signaling.

### SEMA receptors negatively regulate Wnt signaling at the level of CTNNB1

To further investigate the mechanisms of SEMA receptor-mediated Wnt pathway inhibition, we utilized TOPFlash assays in Human Embryonic Kidney (HEK293T) cells, which robustly respond to Wnt pathway activation (Jia et al. 2008). In support of this, Wnt pathway stimulation with ligand (Wnt 3a conditioned media), at the level of the destruction complex with CHIRON, or at the level of CTNNB1 with *CTNNB1^S33Y^*, results in robust Wnt pathway activation (Figure 5A-D). Regardless of the mechanism of pathway activation, *Nrp1* and *Nrp2* significantly inhibit Wnt signaling in HEK293T cells (Figure 5A-D). Further, *Plxna1*^Δ*ECD*^ also inhibits Wnt signaling in these cells (Figure S3A-D), suggesting that NRPs and PLXNs repress Wnt pathway activity in both fibroblasts and epithelial cells.

**Figure 5.**
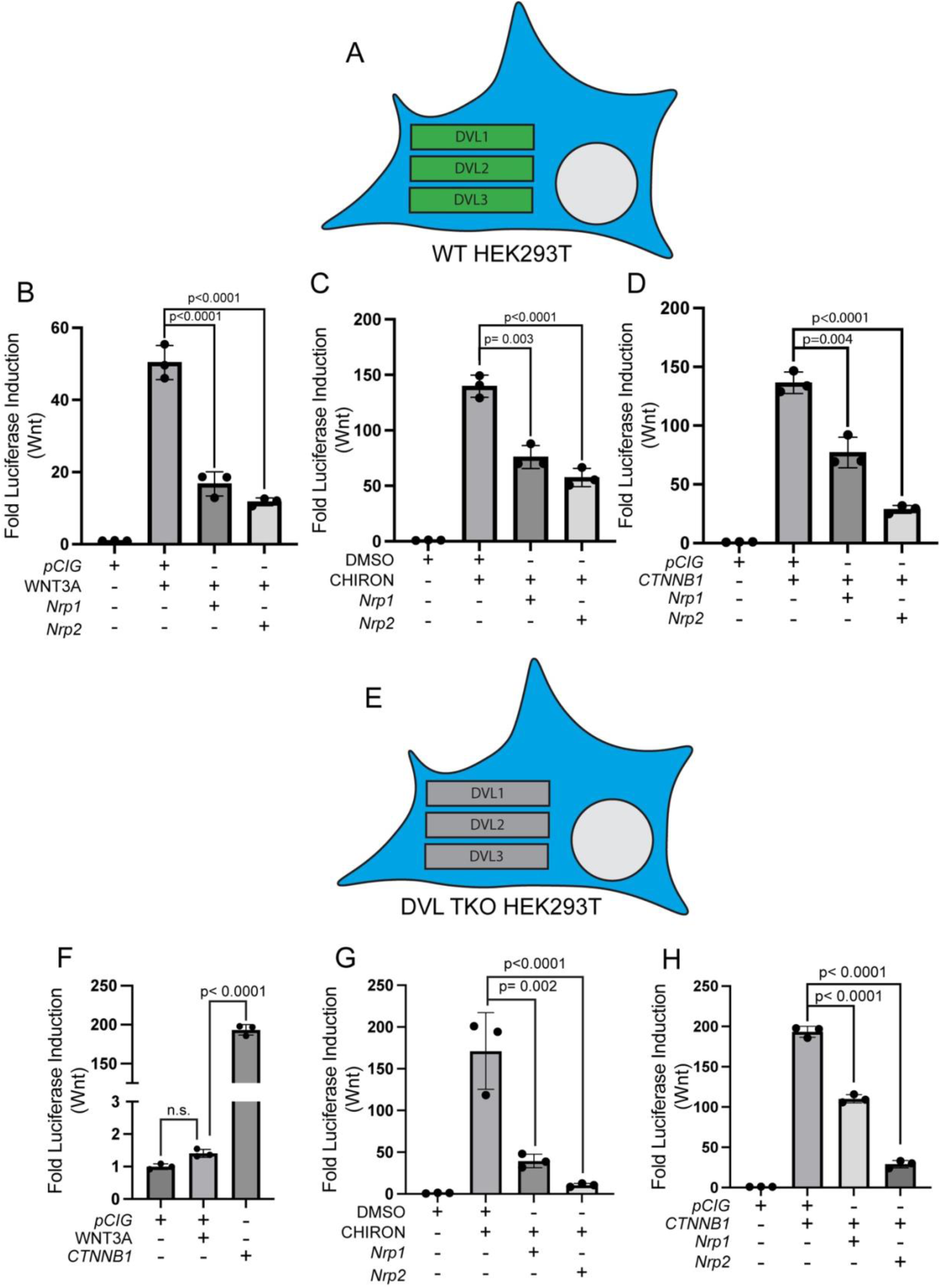
NRPs antagonize Wnt signaling in HEK293T cells in a DVL-independent manner. (**A, E**) Schematic representation of WT or *DVL1^-/-^;DVL2^-/-^;DVL3^-/-^*(DVL TKO) HEK293T cells. (**B-D, F-H**) Wnt-dependent luciferase reporter activity was measured in WT or DVL TKO HEK293T cells transfected with the indicated plasmids and either left untreated or treated with WNT3A conditioned media, DMSO, or CHIRON. Data are representative of at least three biological replicates. Data are reported as mean fold change +/- S.D., with p-values calculated using one-way ANOVA. n.s., not significant.

Dishevelled (DVL) represents a critical link between cell membrane-associated components of the Wnt pathway and the destruction complex, so we next assessed the ability of NRPs and PLXNs to repress Wnt signaling in *DVL1^-/-^,DVL2^-/-^,DVL3^-/-^* HEK293T cells (referred to hereafter as DVL TKO cells) (Klingensmith et al. 1994; Krasnow et al. 1995; Gammons et al. 2016). As expected, DVL TKO cells fail to respond to WNT ligand, although they do respond to CHIRON treatment and *CTNNB1^S33Y^* expression (Figure 5E-H). Importantly, we found that *Nrp1* and *Nrp2* both repress Wnt signaling in DVL TKO cells following stimulation with either CHIRON or *CTNNB1^S33Y^* (Figure 5G-H). *Plxna1*^Δ*ECD*^ also inhibits Wnt signaling in DVL TKO cells (Figure S3E-G). These experiments suggest that SEMA receptors (both NRPs and PLXNs) antagonize Wnt signaling independently of DVL.

### SEMA receptors repress Wnt signaling by CTNNB1 degradation

Since *Nrp1* expression in NIH/3T3 cells reduces endogenous nuclear CTNNB1 (Figure 2I-C’), we sought to assess the effect of SEMA receptors on expression of stabilized CTNNB1 (CTNNB^S33Y^) by western blot analysis of NIH/3T3 cell lysates. While *CTNNB11^S33Y^* expression in NIH/3T3 cells led to significant accumulation of FLAG-tagged CTNNB1, co-expression with *Plxna1* resulted in a greater than 50% decreases in FLAG::CTNNB1^S33Y^ (Figure 6A-B). Co-expression of *CTNNB11^S33Y^* with *Plxna1*^Δ*ECD*^ results in even less CTNNB1^S33Y^, while co-expression with *Plxna1*^Δ*CD*^(which does not antagonize Wnt signaling, cf. Figure 2D), does not alter CTNNB1^S33Y^ levels (Figure 6A-B). To confirm that decreased CTNNB1^S33Y^ protein was not a consequence of altered transcription, we analyzed *FLAG::CTNNB1^S33Y^* expression by qRT-PCR and found that *FLAG::CTNNB1^S33Y^* expression was unaffected by co-expression with *Plxna1*^Δ*ECD*^ (Figure 6C).

**Figure 6.**
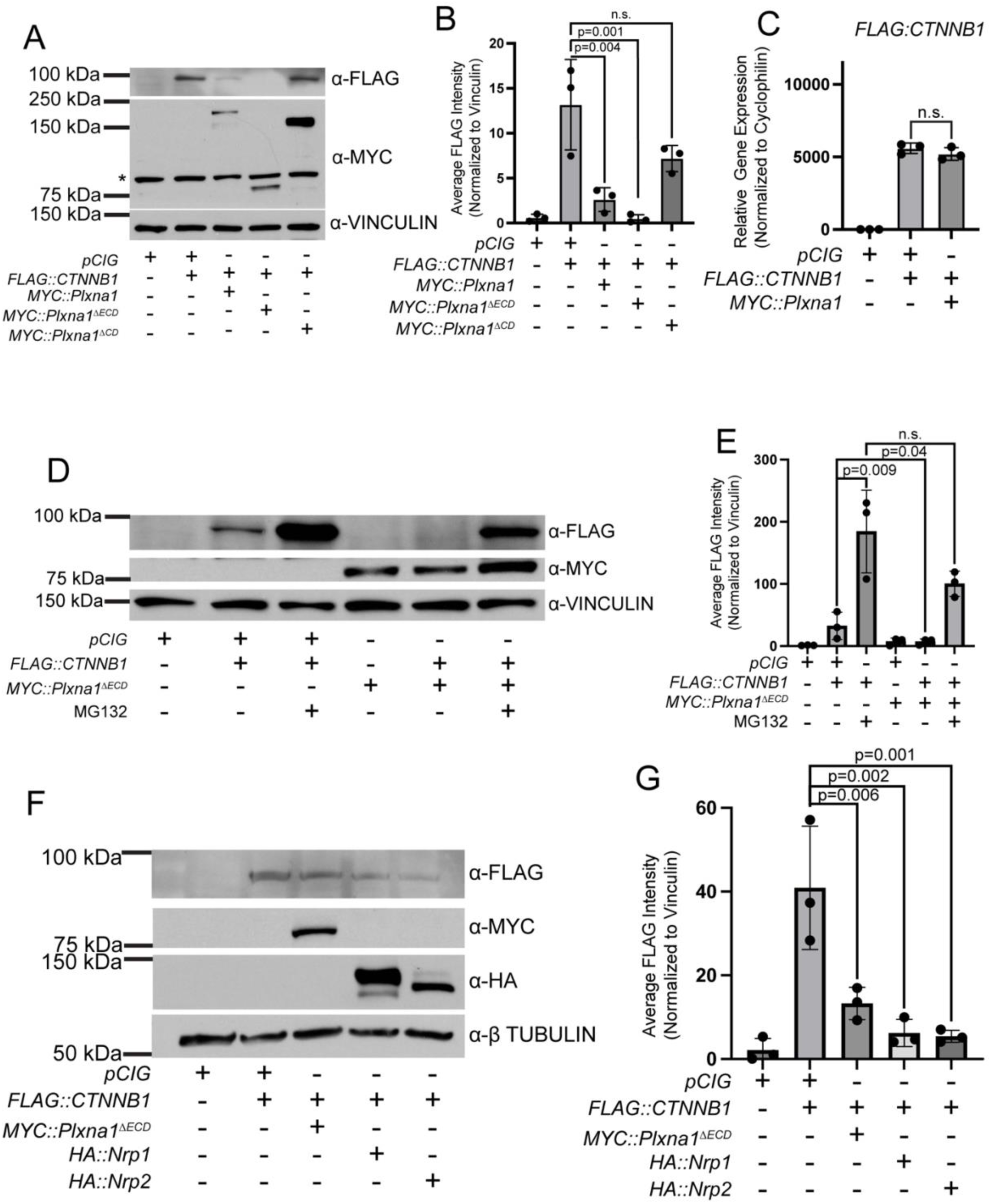
SEMA receptors antagonize Wnt signaling through CTNNB1 degradation. (**A**) Western blot analysis of FLAG::CTNNB1^S33Y^ protein following transfection of NIH/3T3 cells with the indicated plasmids. MYC staining indicates MYC::PLXN protein expression. Asterisk (*) indicates a non-specific band present in untransfected cells. VINCULIN staining was used as a loading control. (**B**) Quantitation of FLAG::CTNNB1^S33Y^ levels in (**A**) normalized to VINCULIN. (**C**) qRT-PCR analysis of *FLAG::CTNNB1^S33Y^* expression following transfection of NIH/3T3 cells with the indicated plasmids. Data points indicate technical replicates. Fold changes were determined using the ΔΔCT method normalized to *Cyclophilin* expression. Data are representative of at least three biological replicates. Data are reported as mean fold change +/- S.D., with p-values calculated using two-tailed Student’s t-test. n.s., not significant. (**D**) Western blot analysis of FLAG::CTNNB1^S33Y^ protein following transfection of NIH/3T3 cells with the indicated plasmids. MYC staining indicates MYC::PLXNA1^ΔECD^ protein expression. Cells were treated with MG132 or vehicle (DMSO) for 24 hours prior to harvesting, as indicated. (**E**) Quantitation of FLAG::CTNNB1^S33Y^ levels in **D** normalized to VINCULIN. (**F**) Western blot analysis of FLAG::CTNNB1^S33Y^ protein following transfection of HEK293T cells with the indicated plasmids. MYC staining indicates MYC::PLXN^ΔECD^ protein expression. HA staining indicates HA:NRP protein expression. (**G**) Quantitation of FLAG::CTNNB1^S33Y^ levels in (**F**) normalized to VINCULIN. Western blot quantitation is reported as mean fold change +/- S.D., with p-values calculated using one-way ANOVA. n.s., not significant. Data are representative of at least three biological replicates.

PLXN- and NRP-mediated destabilization of stabilized CTNNB1 protein was a particularly surprising result. In search of a potential mechanism, we investigated the role of the proteasome in the regulation of CTNNB1^S33Y^ levels. Strikingly, co-expression of *CTNNB1^S33Y^* and *Plxna1*^Δ*ECD*^ in NIH/3T3 cells followed by proteasomal inhibition by MG132 treatment significantly restored CTNNB1^S33Y^ levels (Figure 6D-E). These data suggest that PLXNs and NRPs antagonize Wnt signaling through proteosome-mediated CTNNB1 degradation. Importantly, co-expression of *CTNNB1^S33Y^* with either *Plxna1*^Δ*ECD*^, *Nrp1*, or *Nrp2* in HEK293T cells also reduced levels of CTNNB1^S33Y^ levels (Figure 6F-G).

### NRP-mediated Wnt antagonism requires GSK3β/CK1 phosphorylation of CTNNB1

Canonical regulation of CTNNB1 by the destruction complex is dependent on initial phosphorylation of S45 by CK1 and subsequent phosphorylation of T41, S37, and S33 by GSK3β (Figure 7A) (Amit et al. 2002; Liu et al. 2002). Since CTNNB1 levels are regulated by SEMA receptors through proteasomal degradation, we sought to further assess potential roles of these different phosphorylation sites on PLXN- and NRP-mediated CTNNB1 degradation.

**Figure 7.**
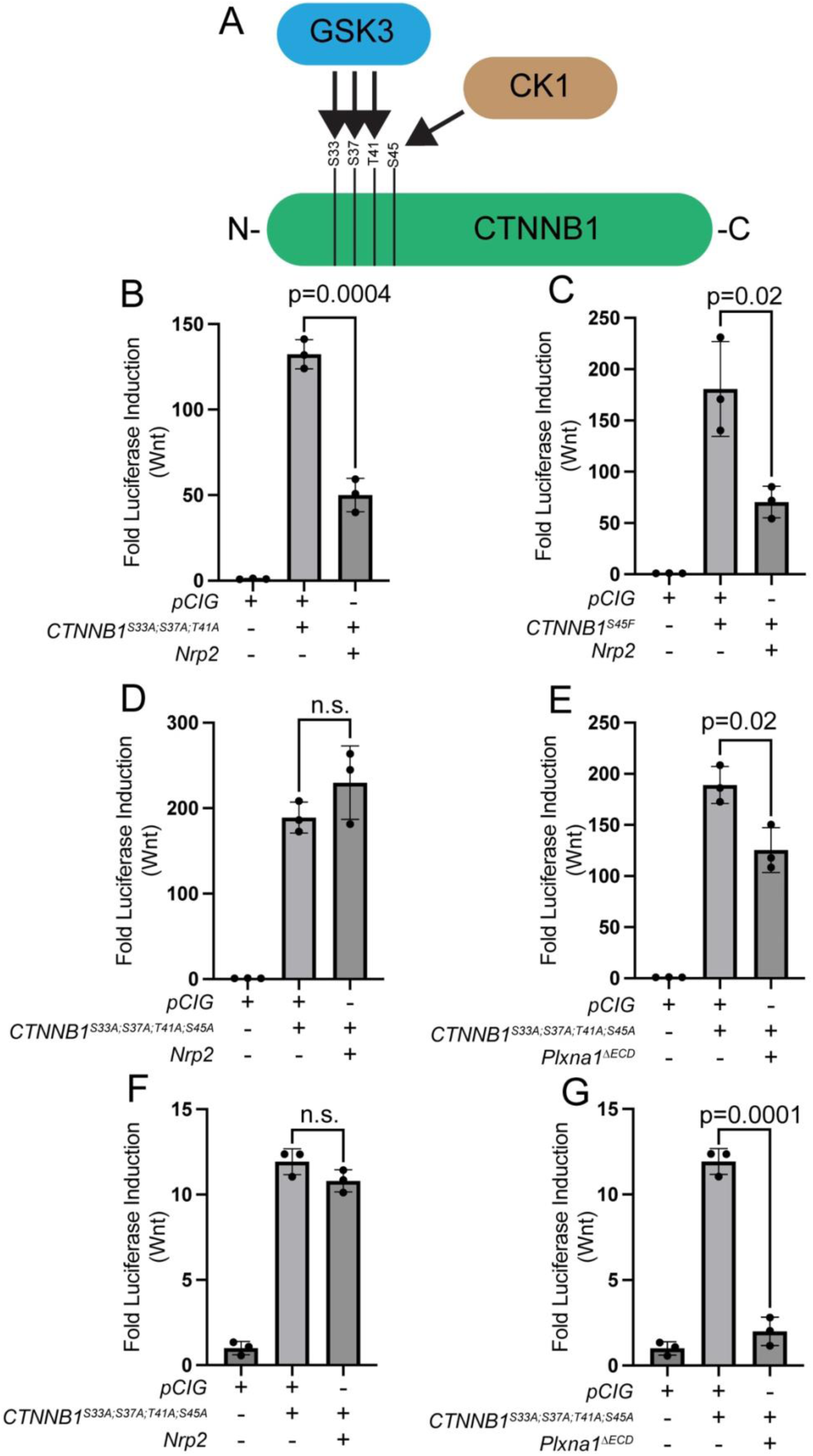
NRPs, but not PLXNs, inhibit Wnt signaling in a GSK3B/CK1-dependent manner. (**A**) Schematic representation of CTNNB1 phosphorylation by CK1 and GSK3B as part of the destruction complex. (**B-E**) HEK293T cells were transfected with the indicated plasmids and Wnt signaling levels were measured by luciferase assay. (**F-G**) NIH/3T3 cells were transfected with the indicated plasmids and Wnt signaling levels were measured by luciferase assay. Data are representative of at least three biological replicates. Data are reported as mean fold change +/- S.D., with p-values calculated using two-tailed Student’s t-test. n.s., not significant.

Individual mutation of either S33, S37, T41 (GSK3β phosphorylation sites) to Alanine does not impact NRP-mediated Wnt pathway antagonism in HEK293T cells (Figure S4A-C). Further, the combined mutation of all of three residues also does not alter NRP-mediated repression of Wnt signaling (Figure 7B). NRP-mediated Wnt antagonism is also unchanged in HEK393T cells expressing *CTNNB1^S45F^*, a non-phosphorylatable mutation in the residue targeted by CK1 (Figure 7C). Strikingly, mutation of all four phosphorylation sites to Alanine does abrogate NRP-mediated Wnt pathway antagonism in HEK293T cells (Figure 7D). Notably, this contrasts with PLXN-mediated Wnt repression, which is unaltered in HEK293T cells expressing *CTNNB1* constructs with four mutated phosphorylation sites (Figure 7E). Importantly, similar results are obtained in NIH/3T3 cells (Figure 7F-G). Together, these data suggest that, although both PLXNs and NRPs restrict Wnt signaling through CTNNB1 degradation, NRPs employ a GSK3β/CK1-dependent mechanism to achieve this degradation, while PLXNs act independently of GSK3β/CK1 phosphorylation of CTNNB1. To confirm the specificity of SEMA receptor-mediated antagonism of Wnt signaling, we treated NIH/3T3 cells with BMP2 ligand and analyzed BMP pathway activity by measuring *Id1, Id2,* and *Id3* expression by RT-qPCR (Figure S5). BMP pathway levels are unaffected following transfection of NIH/3T3 cells with *Plxna1*^Δ*ECD*^, demonstrating selective effects of SEMA receptors on HH and Wnt signaling (Figure S5A-C).

## Discussion

Wnt signaling plays essential roles in a variety of developmental processes, adult tissue homeostasis, and diseases states; however, we still do not possess a complete understanding of this core developmental signaling pathway. Here, we present evidence that SEMA receptors, PLXNs and NRPS, antagonize the Wnt pathway in both fibroblasts and epithelial cells. Further, we find that NRPs and PLXNs repress Wnt signaling independently of primary cilia, and independently of DVL, instead acting at the level of CTNNB1 stability (Figure 8). Notably, our data indicate distinct mechanisms of NRP- (GSK3β/CK1-dependent) and PLXN- (GSK3β/CK1-independent) mediated CTNNB1 degradation. Taken together, these data identify PLXNs and NRPs as novel negative regulators of the Wnt signaling pathway.

**Figure 8.**
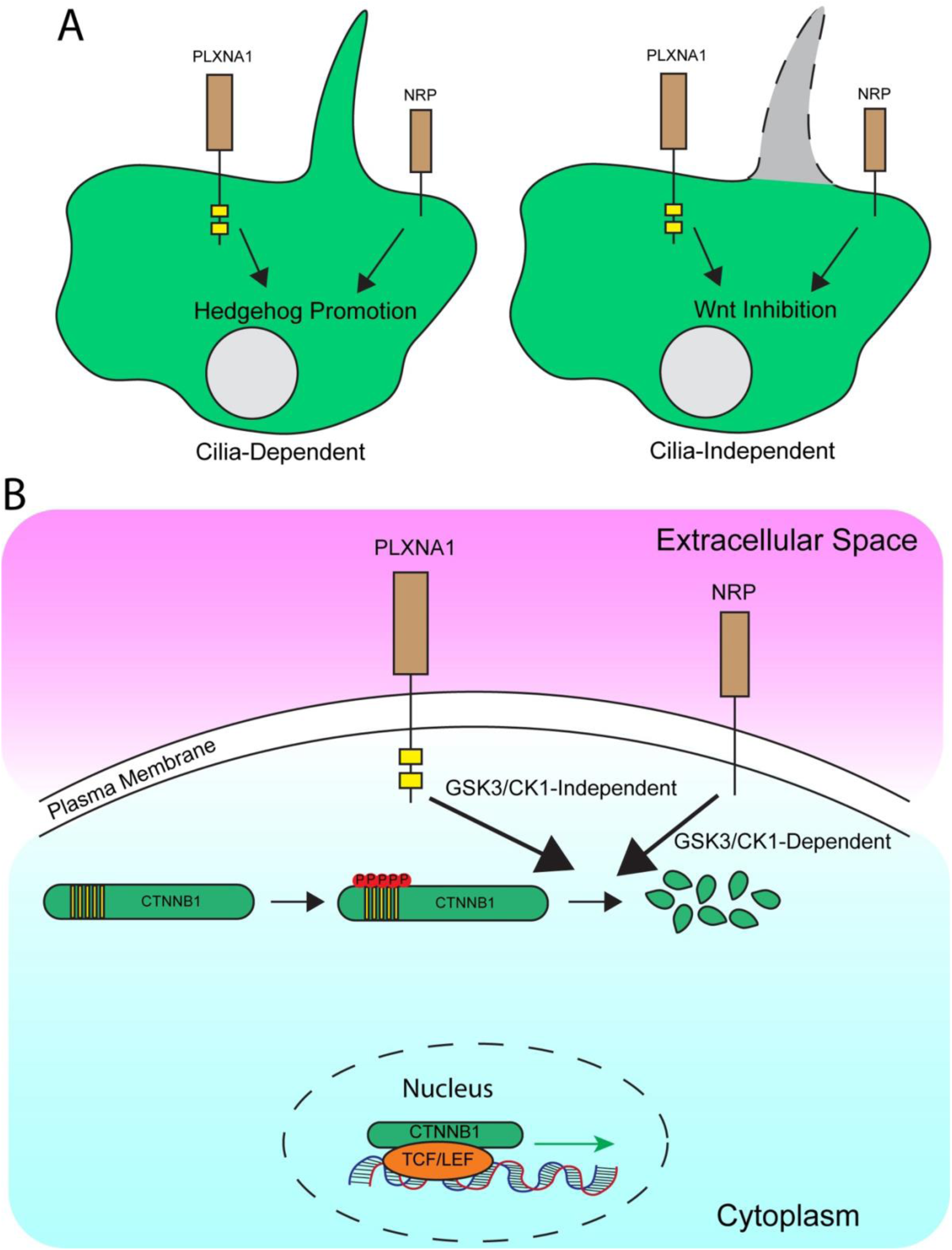
Proposed model of SEMA receptor regulation of HH and Wnt signaling. (**A**) PLXNs and NRPs promote HH signaling (left) in a primary cilia-dependent manner but inhibit Wnt signaling (right) in a primary cilia-independent manner. (**B**) PLXNs and NRPs repress Wnt signaling via distinct mechanism, with NRP-mediated Wnt antagonism dependent on both CK1 and GSK3 activity, while PLXN-mediated Wnt antagonism is independent of CK1 and GSK3. However, both PLXNs and NRPs promote CTNNB1degradation in a proteasome-dependent fashion.

### SEMA receptors as regulators of multiple signaling pathways

Although NRPs and PLXNs were originally described as receptors for SEMA ligands important for axon guidance, both proteins have been implicated in a variety of other signaling pathways, processes, and interactions (He and Tessier-Lavigne 1997; Kolodkin et al. 1997; Takahashi et al. 1999; Tamagnone et al. 1999). NRPs and PLXNs play significant roles in HH signaling (Hillman et al. 2011; Ge et al. 2015; Pinskey et al. 2017; Pinskey et al. 2022). NRPs and PLXNs also contribute to immune system activation, where they are important for T-cell and dendritic cell function (Romeo et al. 2002; Tordjman et al. 2002; Walzer et al. 2005). NRPs and PLXNs also contribute to angiogenesis and vascular development through vascular epithelial growth factor (VEGF) and NF-κβ signaling (Neufeld et al. 2002; Catalano et al. 2009; Glinka et al. 2012; Neufeld et al. 2012). Additionally, NRPs and PLXNs also work with Integrins to modulate cell signaling and cell-cell and cell-matrix interactions (Barberis et al. 2004; Zheng et al. 2009; Goel and Mercurio 2012; Goel et al. 2013).

A multitude of binding partners have been identified for NRPs and PLXNs. NRPs share an affinity for Transforming Growth Factor beta-1 (TGF-β1) and its receptors, Epidermal Growth Factor (EGF), Hepatocyte Growth Factor (HGF), and Platelet-Derived Growth Factor (PDGF) (Yaramis et al. 1998; Parikh et al. 2003; West et al. 2005; Glinka and Prud’homme 2008; Muhl et al. 2017). PLXNs associate with a variety of signaling proteins, including Off-track, RON, MET, and Rho-like GTPases (Rohm et al. 2000; Whitford and Ghosh 2001; Giordano et al. 2002; Conrotto et al. 2004). These established contributions to multiple pathways combined with the broad expression of PLXNs and NRPs in developing and adult tissues, suggest that they play multi-functional roles in the same tissue (Kitsukawa et al. 1995; Perala et al. 2005).

### Mechanisms of PLXN and NRP function in Wnt pathway antagonism

In their traditional role in axon guidance, NRPs and PLXNs form a co-receptor complex to recognize Class 3 Semaphorin ligands (Tamagnone et al. 1999). In other contexts, however, NRPs and PLXNs function independently. Indeed, we find here that PLXN antagonize Wnt signaling in a NRP-independent fashion. Regardless, both NRPs and PLXNs converge at the level of CTNNB1 regulation. An outstanding question is how NRPs and PLXNs regulate CTNNB1 levels.

The PLXN intracellular domain is required for Wnt, pathway antagonism, similar to the requirement for PLXN-mediated HH pathway promotion (Pinskey et al. 2022). Deletion of the PLXN extracellular domain results in exacerbated Wnt inhibition, suggesting that intracellular GAP activity is responsible for PLXN-mediated Wnt pathway antagonism. This could occur through RAP2, a member of the RAS family of GTPases and a modulator of Wnt signaling that is also a substrate of the PLXN GAP domain (Park et al. 2013; Chen et al. 2018). Alternatively, RHO-family GTPases have also been implicated in Wnt planar cell polarity and Wnt-dependent neurite guidance (Marlow et al. 2002; Kishida et al. 2004). ARF6 is a GTPase that regulates N-Cadherin-associated CTNNB1, resulting in its sequestration or liberation and subsequent relocation to the nucleus in melanoma cells (Grossmann et al. 2013). Interestingly, ARF6 is also a substrate of PLXND1 (Sakurai et al. 2010). It is tempting to speculate that PLXNs could also utilize ARF6 to regulate Wnt signaling. PLXN-dependent interactions with these kinases merit further study in the context of Wnt pathway regulation.

While NRPs antagonize Wnt signaling through GSK3β/CK1-dependent CTNNB1 degradation, the means by which NRP achieves this outcome remains to be elucidated. Given the promiscuous binding of NRPs, it is possible that NRP-dependent Wnt antagonism is achieved through a binding partner. For example, NRP interactions with VEGF or VEGFR could modulate Wnt signaling levels. In support of this, previous studies have identified Wnt pathway components as VEGF signaling targets (Li et al. 2021). Alternatively, NRP-mediated Wnt pathway inhibition could be achieved through binding partner-dependent regulation of NF-κβ signaling, which antagonizes Wnt signaling (Xiao et al. 2023). Another possible explanation for NRP-mediated Wnt antagonism could be through the Tyrosine Kinase, FER. Previous studies have identified FER as both an interacting partner of NRP1 and a negative regulator of Wnt signaling that functions at the level of CTNNB1 (Jiang et al. 2010; Putzke and Rothman 2010). Future studies should assess the potential dependence of FER kinase on NRP-mediated Wnt repression.

NRP-mediated Wnt repression could depend on upstream modulation of CK1 and GSK3β activity. Indeed, some studies have already identified the NRP/PLXN co-receptor complex as well as NRP2 alone as positive and negative regulators of GSK3β, respectively (Montolio et al. 2009; Ng et al. 2016). However, the link between CK1 and NRPs is less clear. One possibility is Protein Phosphatase 1 (PP-1), which regulates both CK1 and GSK3β in the context of Wnt signaling (Luo et al. 2007). Another possibility is mitogen-activated protein kinase (MAPK), which has been linked to the regulation of both CK1 and GSK3β (Thornton et al. 2008; Knippschild et al. 2014). Notably, MAPK has been identified as a possible regulatory substrate of NRP1 (Wey et al. 2005).

### Consequences of SEMA receptor antagonism of Wnt signaling on organogenesis

The broad expression of NRPs and PLXNs combined with the multitude of Wnt-dependent tissues suggest potential contributions of SEMA receptors to Wnt signaling in both developing and adult tissues. For example, Wnt plays important roles in the developing nervous system, where both PLXNs and NRPs are broadly expressed (Kawakami et al. 1996; Perala et al. 2005; Salinas 2012). Further, Wnt signaling is as an important repulsive axon guidance cue, similarly to the already established role of SEMA signaling (Liu et al. 2005). Similar to neural development, NRPs and PLXNs may influence Wnt signaling during cardiac development and angiogenesis, processes in which both Wnt signaling and SEMA receptors contribute (Parmalee and Kitajewski 2008; Li et al. 2022). Finally, NRPs, PLXNs and Wnt contribute to a number of cancers. Roles for PLXNs and, particularly, NRPs have been described in human colon cancer, a disease where Wnt signaling plays a prominent role (Parikh et al. 2004; Samuel et al. 2011; Bollard et al. 2019; Bai et al. 2020; Zhao et al. 2022). Indeed, a previous study has even identified a correlation between NRP2 and CTNNB1 activity in human gastrointestinal cell lines (Samuel et al. 2011). Future functional studies will be essential to address these possibilities, although technical challenges exist given the number of PLXNs that could play redundant roles in Wnt pathway antagonism. A final consideration will be potential NRP/PLXN-regulated crosstalk between pathways (e.g., HH, Wnt, VEGF) in both normal and disease contexts. Previously established crosstalk between HH and Wnt signaling in the development of colorectal cancer would make this a tempting system to interrogate (Song et al. 2015).

## Acknowledgements

We are grateful to Dr. Stephane Angers (University of Toronto, ON, CA) for providing *DVL* TKO HEK293T cells. We also acknowledge Dr. David Sabatini (Institute of Organic Chemistry and Biochemistry, CZ) and Dr. Kris Wood (Duke University, NC, USA) for providing the *CTNNB1^S33A;S37A;T41A;S45A^* plasmid. Members of the Allen and Cadigan labs contributed technical assistance, comments, and suggestions. In particular, we thank Dr. Richard Stewart for valuable interactions. We are also thankful to Drs. Sue O’Shea and Kristen Verhey for sharing equipment and reagents. Confocal imaging was performed in the Microscopy Core at the University of Michigan Medical School. This study was supported by funding from the National Institutes of Health to BLA (R01 CA275182) and RJG (R01 MH119346).

## Author Contributions

Tyler M Hoard, Conceptualization, Formal analysis, Validation, Investigation, Methodology, Writing-original draft, Writing-review and editing. Katie Liu, Investigation. Ken M Cadigan, Conceptualization, Investigation. Roman J Giger, Conceptualization, Investigation. Ben L Allen, Conceptualization, Resources, Formal analysis, Supervision, Funding acquisition, Investigation, Methodology, Writing-original draft, Project administration, Writing-review and editing.

## Materials and Methods

### Cell culture

Cells were maintained in Dulbecco’s Modified Eagle Medium (DMEM; Thermo Fisher Scientific, 11965-118) supplemented with 10% Bovine Calf Serum (ATCC, 30-2030) and 1X Penicillin-Streptomycin-Glutamine (Life Technologies, 10378016). Cells were incubated at 37°C with 5% CO_2_ and 95% humidity. NIH/3T3 cells (CRL-1658) and HEK293T (CRL-3216) were purchased from ATCC. *DVL1^-/-^;DVL2^-/-^;DVL3^-/-^*(DVL TKO) cells were a generous gift from Dr. Stephane Angers at the University of Toronto (Gammons et al. 2016). *Kif3a^-/-^* NIH/3T3 Flp-In cells were obtained from Dr. Kristen Verhey (Engelke et al. 2019). L Wnt-3a cells for producing WNT3A conditioned media was obtained from ATCC (CRL-2647). All cell lines were mycoplasma negative as confirmed by testing with (Kit details needed).

### Generation of *Nrp1^-/-^;Nrp2^-/-^* NIH/3T3 fibroblasts

A CRISPR Gene Knockout Kit from Synthego was used to generate *Nrp1^-/-^;Nrp2^-/-^*NIH/3T3 fibroblasts. 2.5 ×10^4^ NIH/3T3 fibroblasts were reverse-transfected using Lipofectamine CRISPRMAX Cas9 transfection reagents (Invitrogen, CMAX00008) with 0.13 μM sgRNA and 0.1 μM spCas9 2NLS nuclease and plated in 500 μL in a 24-well plate. Cells were split 48h later to generate single-cell colonies. Upon reaching 40-60% confluency, monoclonal colonies were split into 6-well plates and grown to 60% confluency. Cells were then lysed using QuickExtract DNA Extraction Solution (BioSearch Technologies, QE0905T) following the manufacturer’s instructions for genotyping PCR. Genotyping primer, sequencing primer, and sgRNA sequences are included below (5’-3’):

*mNrp1* F: CCCGCTGAGGATTTTCTGGT

*mNrp1* R: CAGGAGAAGCCAGCAACCAA

*mNrp1* Sequencing: TTAAGAGCGTTTCGGATTGTTAAGATTATC

*mNrp1* sgRNA1: GCGAGCGUGGCGCACAGCAA

*mNrp1* sgRNA2: CCGGCUGUCACUUACCGCUG

*mNrp1* sgRNA3: CCCCUUCGCCCGAGGGGACU

*mNrp2* F: TGTCTTGGGAGGGGGAGTAG

*mNrp2* R: TGTTTAAGTGGCCCTCTGTGG

*mNrp2* Sequencing: GTAGGAGGAGAGGAGAGTTGAAATAGTCTC

*mNrp2* sgRNA1: CGACCUCCGCAGGGUGGAUC

*mNrp2* sgRNA2: AUAGUCCUGGGGGUAGCCUG

*mNrp2* sgRNA3: AAUCUUCUGGUUGGGUUCGG

### Generation of mutant *CTNNB1* constructs

Individual and combination mutant versions of *Homo sapiens CTNNB1* DNA constructs were generated using standard cloning techniques and the QuickChange II XL Site-Directed Mutagenesis Kit (Agilent Technologies, 200521), unless otherwise noted. *CTNNB1^S45F^* was a generous gift from Dr. Ken Cadigan. *CTNNB11^S33A;S37A;T41A;S45A^-pcw107-V5* was originally obtained as a gift from Drs. David Sabatini and Kris Wood (Addgene plasmid #64613; http://n2t.net/addgene:64613; RRID: Addgene_64613) (Martz et al. 2014).

*CTNNB1^S33A;S37A;T41A;S45A^* was then cloned into the pCIG vector, which contains a CMV enhancer, chicken-beta-actin promoter, and internal ribosome entry site (IRES) with a nuclear enhanced green fluorescent protein reporter (3XNLS-EGFP) (Megason and McMahon 2002).

### Cell signaling assays

Luciferase-based reporter assays in NIH/3T3 and HEK293T cells were performed as previously described using a ptcΔ136-GL3 reporter construct to measure HH activity or TOP-FLASH for Wnt activity (Molenaar et al. 1996; Nybakken et al. 2005). Briefly, NIH/3T3 cells were seeded at a 5×10^4^ cells per well and HEK293T cells were seeded at 1×10^5^ cells per well into 0.5% gelatin-coated 24-well plates. The following day, cells were transfected with empty vector (pCIG) or experimental constructs along with ptcΔ136-GL3 or TOP-FLASH luciferase reporter constructs and β-galactosidase transfection control (pSV-β-galactosidase; Promega, E1081). Transfections were performed using Lipofectamine 2000 (Invitrogen, 11668) and Opti-MEM reduced serum media (Invitrogen, 31985). Then, 24 h after transfection, culture media were replaced with fresh 10% media containing DMSO, 300 nM SAG (Enzo Life Sciences, ALX-270-426-M001), 30 μM CHIRON (APExBIO, A3011), or 50% WNT3A conditioned media (CM). WNT3A CM was produced by growing L-Wnt3a cells in 20 cm plates until confluent. After cells reached confluency, media was replaced with 10mL of fresh media. CM was collected and sterile filtered 48 h later. Luciferase reporter activity and Beta-Galactosidase activity were measured 24 h later on a Spectramax M5^e^ Plate Reader (Molecular Devices) using the Luciferase Assay System (Promega, E1501) and the Betafluor Beta Galactosidase Assay Kit (EMD Millipore, 70979), respectively. Luciferase values were divided by beta-galactosidase values to control for transfection efficiency, and the data were reported as fold induction relative to the vector-transfected control. Treatments were performed in triplicate, with each datapoint representing a technical replicate, and averaged (bar height), with error bars representing the standard deviation between triplicates. Each experiment was repeated for at least three biological replicates, and representative results are shown. For comparison of two data sets, student’s unpaired t-test was used to determine whether each treatment was significantly different from the control. For comparisons between more than two data sets, a one-way ANOVA was used. A p-values of 0.05 was designated as significant.

### Immunofluorescent analysis of cultured cells

NIH/3T3 fibroblasts were plated at 1.5×10^5^ cells per well onto glass coverslips in 6-well dishes. Cells were transfected 24 h after plating using Lipofectamine 2000 (Invitrogen, 11668) and Opti-MEM reduced serum medium (Invitrogen, 31985). Then, 24 h after transfection, media were replaced by fresh media containing DMSO or 30 μM CHIRON (APExBIO, A3011). After 24 h, cells were washed with PBS and fixed in 4% for 15 min at room temperature and washed with PBS. Cells were then fixed with ice cold methanol at −20°C for 5 min. After being washed with PBS, cells were permeabilized for 5 min at room temperature with 0.2% Triton X-100 in PBS prior to staining. Primary antibodies are included in **Table 1** and were diluted in IF blocking buffer (30 g/L bovine serum albumin, 1% heat-inactivated sheep serum, 0.02% NaN_3_, and 0.1% Triton X-100 in PBS). Coverslips were incubated with primary antibodies for 1 h at room temperature, followed by a 10 min DAPI stain (1:30,000 in 1X PBS at room temperature), and 1 h incubation at room temperature with secondary antibodies included **Table 2**. Coverslips were mounted to glass slides using Shandon Immu-Mount Mounting Medium (Fisher, 9990412).

**Table 1.** A list of primary antibodies used in experiments.

| Primary Antibody (Species) | Dilution | Source | Identifier |
| --- | --- | --- | --- |
| HA (Rabbit IgG) | 1:10,000 | Bethyl Labs | A190-108A |
| CTNNB1 (Mouse IgG1) | 1:200 | BD Biosciences | 610153 |
| β-Tubulin (Mouse IgG1) | 1:10,000 | Developmental Studies Hybridoma Bank | E7 |
| NRP1 (Goat IgG) | 1:200 (IF)<br>1:2,000 (WB) | R&D Systems | AF566 |
| NRP2 (Rabbit IgG) | 1:500 | Cell Signaling | 3366S |
| FLAG (Mouse IgG1) | 1:500 | Sigma | F3165 |
| MYC (Rabbit IgG) | 1:5,000 | Bethyl Labs | A190-105A |
| HA (Mouse IgG2b) | 1:1,000 (IF) | OriGene | AM00198PU-N |
| HA (Rabbit IgG) | 1:10,000 (WB) | Bethyl Labs | A190-108A |
| VINCULIN (Rabbit IgG) | 1:1000 | Cell Signaling | 13901 |

**Table 2.** A list of secondary antibodies used in experiments.

| Secondary Antibody (species) | Dilution | Source | Identifier |
| --- | --- | --- | --- |
| Alexa555 (Goat anti-Mouse IgG1) | 1:500 | Invitrogen | A21127 |
| Alexa647 (Goat anti-Mouse IgG2b) | 1:500 | Invitrogen | A21242 |
| Alkaline-Phosphatase (Rabbit anti-Mouse IgG) | 1:10,000 | Jackson ImmunoResearch | 211052171 |
| Alkaline-Phosphatase (Goat anti-Donkey IgG) | 1:10,000 | Jackson ImmunoResearch | 705035003 |
| Alkaline-Phosphatase (Mouse anti-Donkey IgG) | 1:10,000 | Jackson ImmunoResearch | 715035150 |

Immunofluorescent analysis and imaging were performed on a Leica SP5X Upright 2-Photon Confocal microscope using LAS AF software (Leica) and a Leica 63X (type: HC Plan Apochromat CS2; NA 1.2) water immersion objective. Nuclear CTNNB1 was quantified using ImageJ, with signal intensity being normalized to signal quantitation in an area in the cell adjacent to the nucleus. Average values are represented by bar height, and error bars represent standard deviation among samples. Student’s *t*-tests were used compare statistical significance to the control, with p-values of 0.05 or less considered significant.

### Western blot analysis

NIH/3T3 or HEK293T cells were plated at 1.5×10^5^ cells per well in 6-well dishes and transfected 24 h after plating using Lipofectamine 2000 (Invitrogen, 11668) and Opti-MEM reduced serum medium (Invitrogen, 31985). Then, 24 h after transfection, media were replaced by fresh media containing DMSO or 30 μM CHIRON (APExBIO, A3011). Then 24 h later, cells were lysed in radioimmunoprecipitation assay (RIPA) buffer (50 mM Tris-HCL, pH 7.2, 150 mM NaCl, 0.1% Triton X-100, 1% sodium deoxycholate, and 5 mM EDTA). Lysates were centrifuged at 14,000xg for 30 min at 4°C to pellet insoluble components. Protein concentrations were determined using a BCA Protein Assay Kit (Fisher, PI23225). After boiling for 10 min, 50 μg of protein from each sample were separated using SDS-PAGE with 7.5% gels and transferred onto Immun-Blot PVDF membranes (Bio-Rad, 162-0177). Membranes were washed in Tris-buffered saline (TBS) containing 0.5% OmniPur Tween-20 (TBST) and blocked in Western blot blocking buffer (30 g/L bovine serum albumin with 0.2% NaN_3_ in TBST) for 1 h. Antibodies used for staining blots are listed in **Table 1**. Blots were stained with primary antibodies overnight at 4°C and with secondary antibodies (shown in **Table** 2) at room temperature for 1 h. Cytiva Amersham ECL Prime Western Blot Detection Reagent (Fisher, 45-002-401) was added 5 min before membranes were exposed to ByBlot CL Audoradiography Film (Denville, E3018) and developed using a Konica Minolata SRX-101A Medical Film Processor. Quantitation of western blots was performed using ImageJ. Background integrated density intensity was subtracted from band values in each lane. This value was then normalized to the background-corrected integrated density of the housekeeping protein (VINCULIN or β-TUBULIN).

### RT-qPCR

NIH/3T3 or HEK293T cells were cultured and transfected as described above. RNA was isolated using Trizol (Thermo, 15596026) per manufacturer instructions. cDNA was generated using 2 μg of template RNA and a High-Capacity cDNA Reverse Transcription Kit (Applied Biosystems, 4368813). cDNA was diluted 1:100 and qPCR was performed using PowerUp SYBY Green Master Mix (Applied Biosytems, A25742) on an Applied Biosystems Quantstudio 3 Real-Time PCR System with the following primers (5’-3):

ID1 F: CCAGTGGGTAGAGGGTTTGA

ID1 R AGAAATCCGAGAAGCACGAA

ID2 F: ATCAGCCATTTCACCAGGAG

ID2 R: TCCCCATGGTGGGAATAGTA

ID3 F: ACTCAGCTTAGCCAGGTGGA

ID3 R: GTCAGTGGCAAAAGCTCCTC

FLAG:CTNNB1 F: ACGACGATGACAAGGACTACA

FLAG:CTNNB1 R: GAGTAGHCCATTGTCCACGCT

---

**Figure S1.**
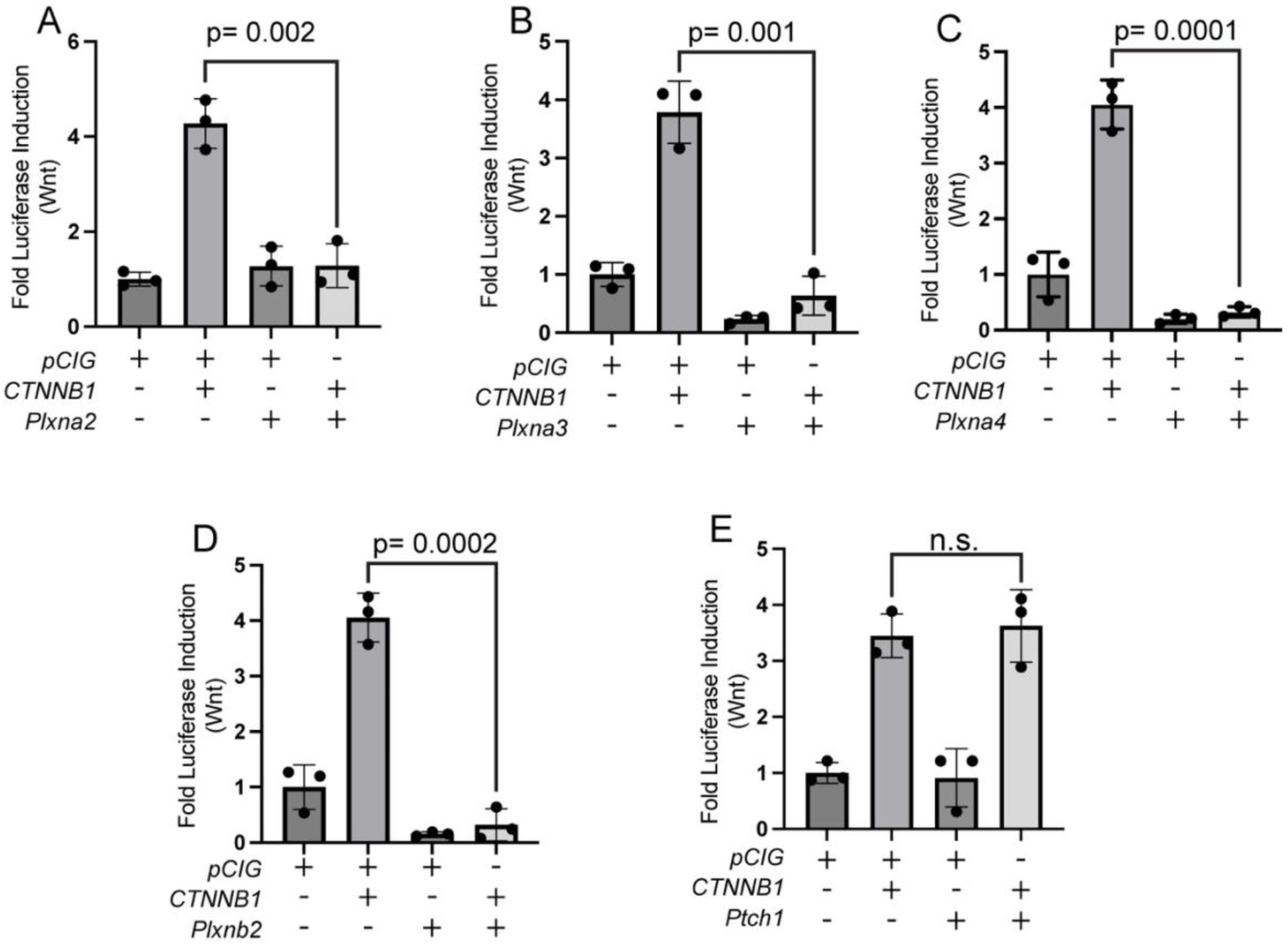
PLXNA and PLXNB subfamily members antagonize Wnt signaling in NIH/3T3 fibroblasts. (**A-E**) Wnt-dependent luciferase reporter activity was measured in NIH/3T3 cells transfected with the indicated plasmids. Data points indicate technical replicates. Data are representative of at least three biological replicates. Data are reported as mean fold change +/- S.D., with p-values calculated using two-tailed Student’s t-test. n.s., not significant.

**Figure S2.**
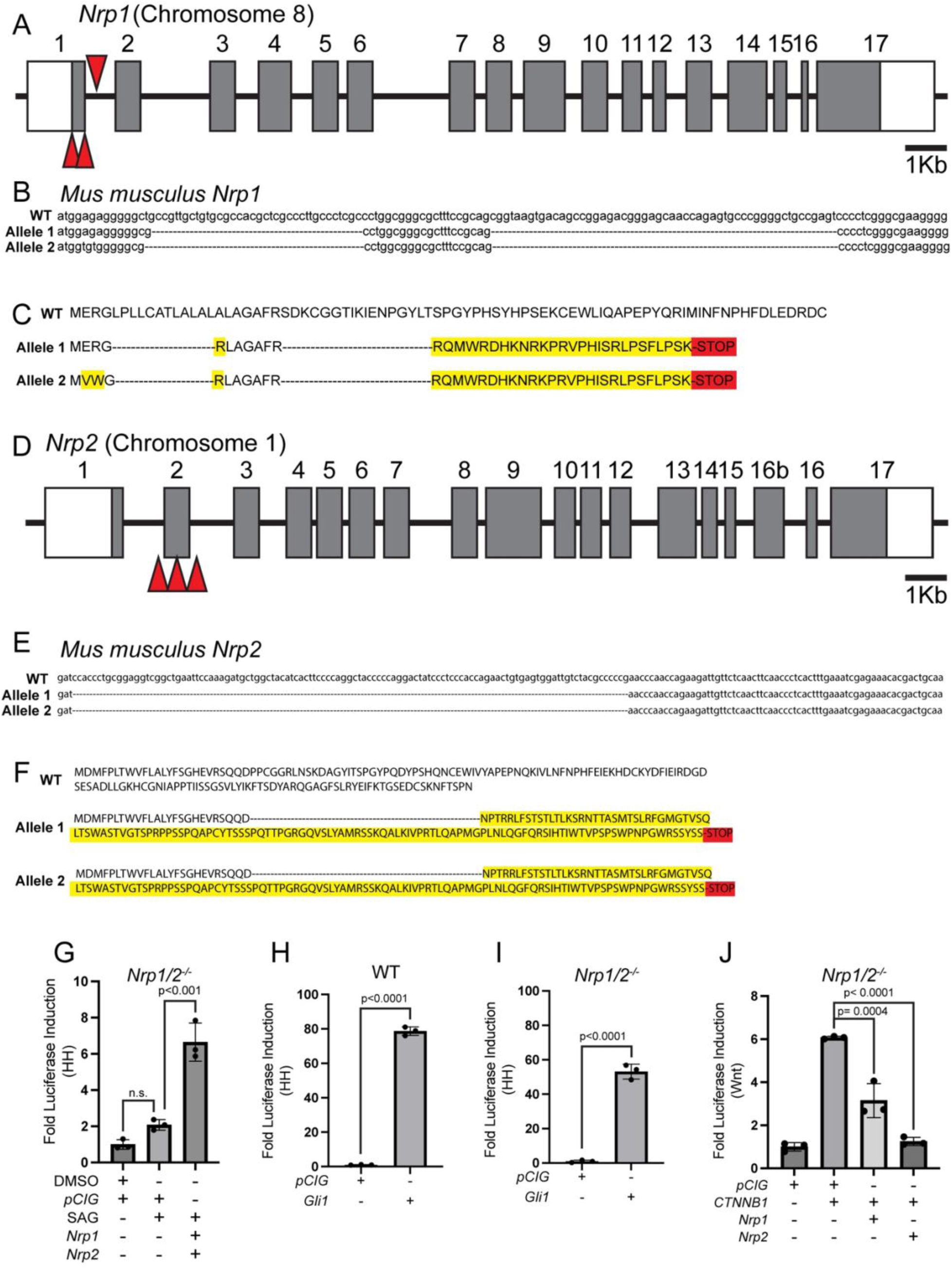
Generation of *Nrp1^-/-^;Nrp2^-/-^* NIH/3T3 cells. (**A,D**) Mouse *Nrp1* (**A**) or *Nrp2* (**D**) locus, including exons (non-coding, white boxes; coding, gray boxes) and introns (lines), with red triangles indicating the target sites of CRISPR gRNAs. Scale bar = 1Kb. (**B, E**) Nucleotide sequence alignment of the WT *Nrp1* (**B**) or *Nrp2* (**E**) sequence and CRISPR-edited alleles. Dashed lines indicate CRISPR-edited nucleotide deletions. (**C, F**) Amino acid sequence alignment of WT and disrupted NRP1 (**C**) or NRP2 (**F**) proteins encoded by the CRISPR-edited *Nrp1* alleles. Dashed lines indicate amino acid deletions; yellow highlighted amino acids represent missense mutations; red highlighting denotes a premature stop codon leading to a truncated protein. (**G**) *Nrp1^-/-^Nrp2^-/-^* NIH/3T3 cells were transfected with the indicated plasmids and HH signaling was measured by luciferase assay. (**H-I**) WT or *Nrp1^-/-^Nrp2^-/-^*NIH/3T3 cells were transfected with the indicated plasmids, and HH signaling was measured by luciferase assay. (**J**) *Nrp1^-/-^Nrp2^-/-^*NIH/3T3 cells were transfected with the indicated plasmids and Wnt signaling levels were measured by luciferase assay. Data are representative of at least three biological replicates. Data are reported as mean fold change +/- S.D., with p-values calculated using two-tailed Student’s t-test. n.s., not significant.

**Figure S3.**
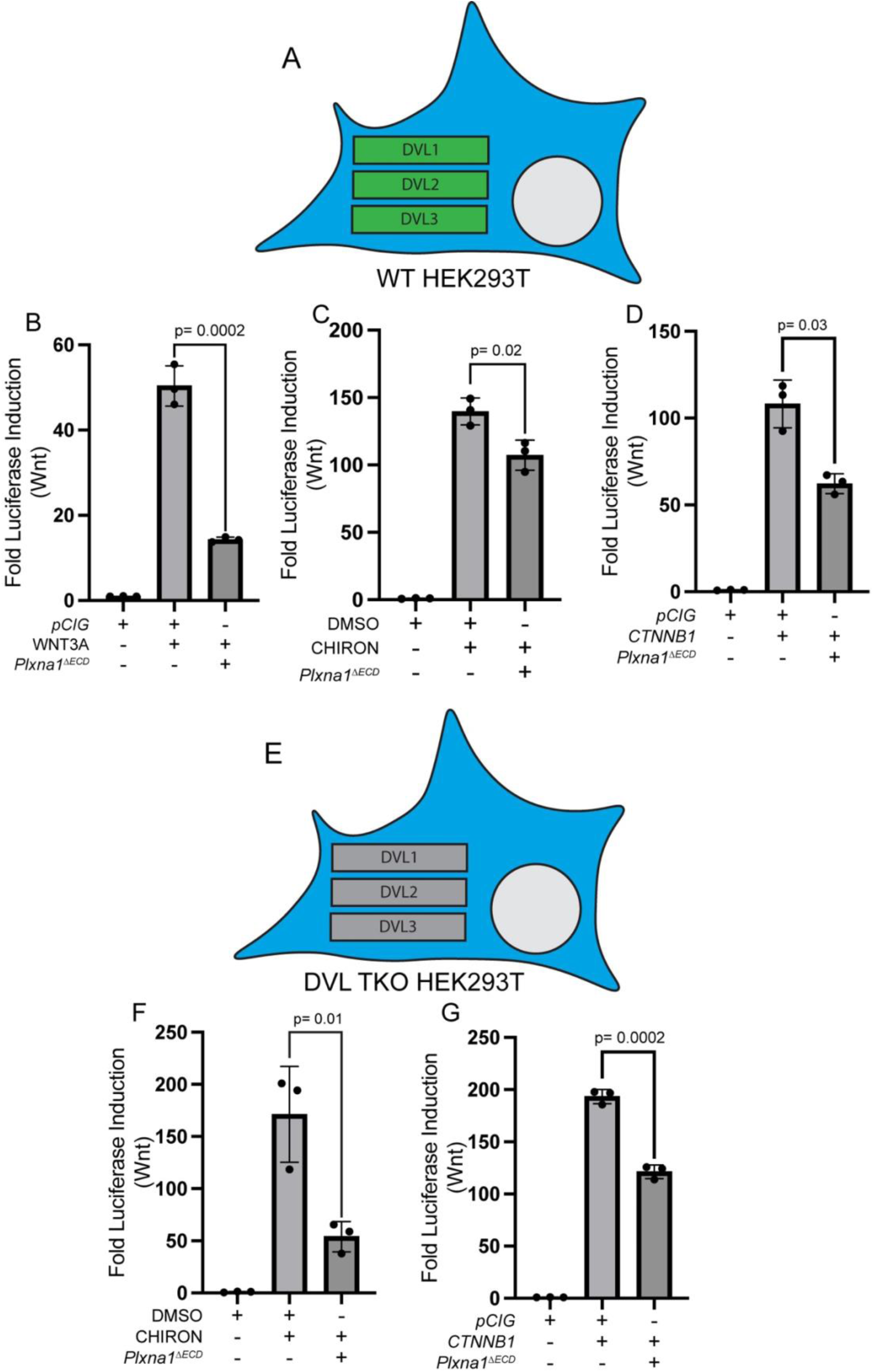
PLXN antagonizes Wnt signaling in HEK293T cells in a DVL-independent manner. (**A, E**) Schematic representation of WT or *DVL^-/-^;DVL2^-/-^;DVL3^-/-^*(DVL TKO) HEK293T cells. (**B-D, F-G**) Wnt-dependent luciferase reporter activity was measured in WT or DVL TKO HEK293T cells transfected with the indicated plasmids and either left untreated or treated with WNT3A conditioned media, DMSO, or CHIRON. Data are representative of at least three biological replicates. Data are reported as mean fold change +/- S.D., with p-values calculated using two-tailed Student’s t-test. n.s., not significant.

**Figure S4.**
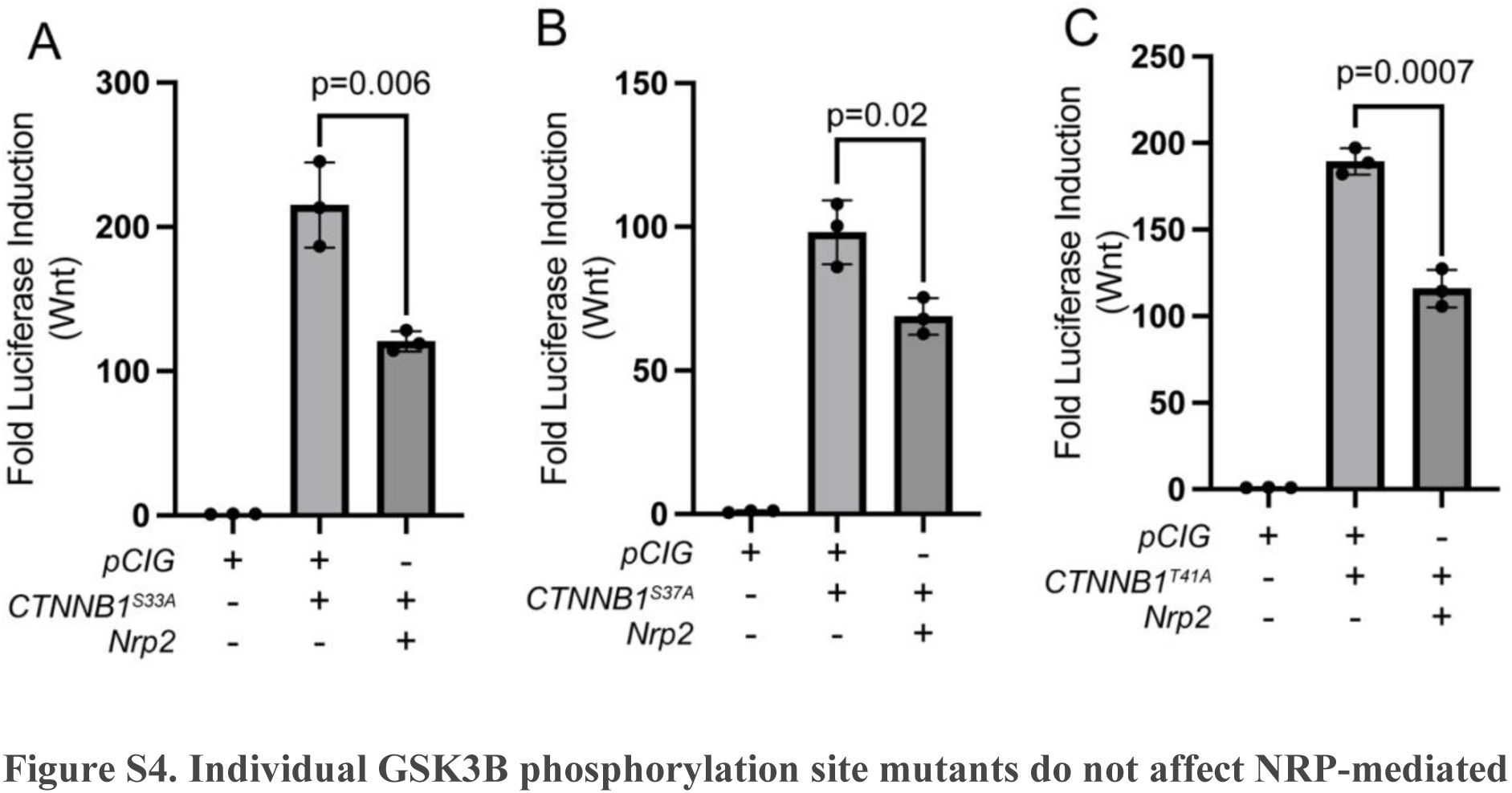
Individual GSK3B phosphorylation site mutants do not affect NRP-mediated Wnt repression. (**A-C**) HEK293T cells were transfected with the indicated plasmids and Wnt signaling levels were measured by luciferase assay. Data are representative of at least three biological replicates. Data are reported as mean fold change +/- S.D., with p-values calculated using two-tailed Student’s t-test. n.s., not significant.

**Figure S5.**
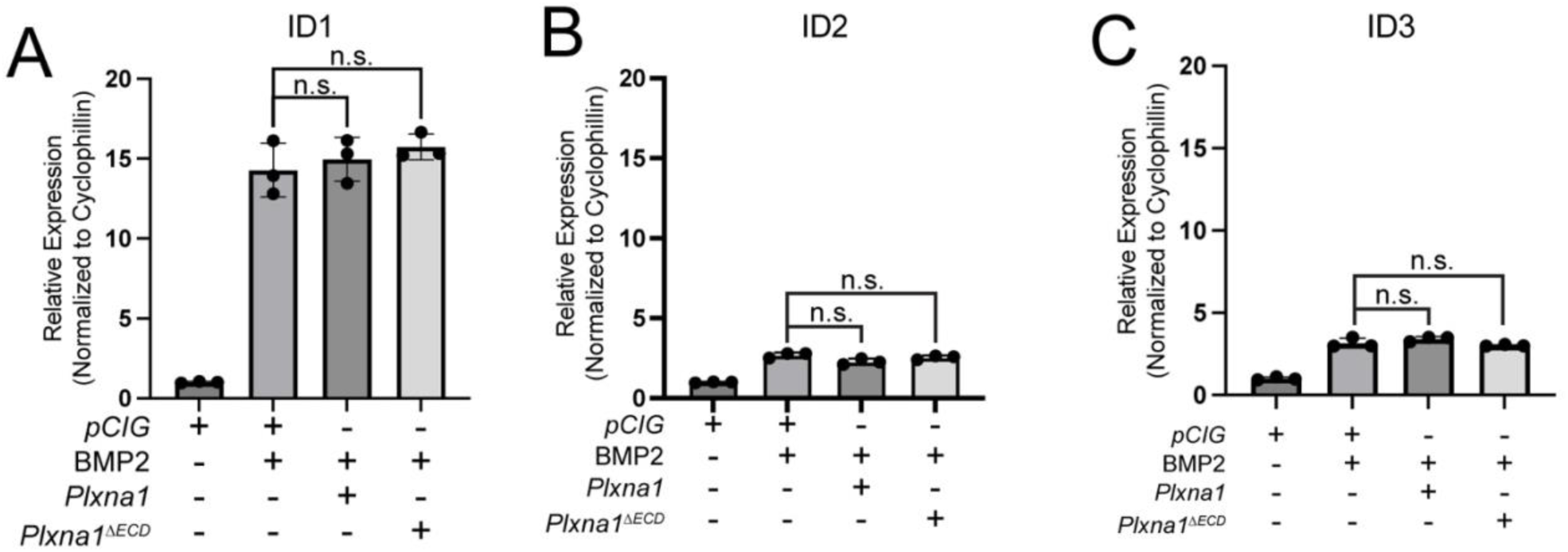
BMP signaling is not modulated by PLXNs. (**A-C**) qRT-PCR analysis of *Id1, Id2,* and *Id3* expression following transfection of NIH/3T3 cells with the indicated plasmids and treatment with BMP2 for 24 hours. Data points indicate technical replicates. Fold changes were determined using the ΔΔCT method normalized to *Cyclophilin*. Data are representative of at least three biological replicates. Data are reported as mean fold change +/- S.D., with p-values calculated using two-tailed Student’s t-test. n.s., not significant.

